# Distinguishing microgliosis and tau deposition in the mouse brain using paramagnetic and diamagnetic susceptibility source separation

**DOI:** 10.1101/2024.04.11.588962

**Authors:** Jayvik Joshi, Minmin Yao, Aaron Kakazu, Yuxiao Ouyang, Wenzhen Duan, Manisha Aggarwal

## Abstract

Tauopathies, including Alzheimer’s disease (AD), are neurodegenerative disorders characterized by hyperphosphorylated tau protein aggregates in the brain. In addition to protein aggregates, microglia-mediated inflammation and iron dyshomeostasis are other pathological features observed in AD and other tauopathies. It is known that these alterations at the subcellular level occur much before the onset of macroscopic tissue atrophy or cognitive deficits. The ability to detect these microstructural changes with MRI therefore has substantive importance for improved characterization of disease pathogenesis. In this study, we demonstrate that quantitative susceptibility mapping (QSM) with paramagnetic and diamagnetic susceptibility source separation has the potential to distinguish neuropathological alterations in a transgenic mouse model of tauopathy. 3D multi-echo gradient echo data were acquired from fixed brains of PS19 (Tau) transgenic mice and age-matched wild-type (WT) mice (n = 5 each) at 11.7 T. The multi-echo data were fit to a 3-pool complex signal model to derive maps of paramagnetic component susceptibility (PCS) and diamagnetic component susceptibility (DCS). Group-averaged signal fraction and composite susceptibility maps showed significant region-specific differences between the WT and Tau mouse brains. Significant bilateral increases in PCS and |DCS| were observed in specific hippocampal and cortical sub-regions of the Tau mice relative to WT controls. Comparison with immunohistological staining for microglia (Iba1) and phosphorylated-tau (AT8) further indicated that the PCS and DCS differences corresponded to regional microgliosis and tau deposition in the PS19 mouse brains, respectively. The results demonstrate that quantitative susceptibility source separation may provide sensitive imaging markers to detect distinct pathological alterations in tauopathies.

## 1. INTRODUCTION

Tauopathies, including Alzheimer’s disease (AD), are neurodegenerative disorders characterized by abnormal intracellular accumulation of hyperphosphorylated tau protein aggregates in the brain. In AD, a secondary tauopathy, the spread of tau neurofibrillary tangles has been shown to be more closely correlated with local neurodegeneration and cognitive decline compared to amyloid-β deposits (Arriagada et al., 1992; Bejanin et al., 2017). In addition to the formation of protein aggregates, microglia-mediated neuroinflammation and iron dyshomeostasis are other key pathological features observed in both human (Kenkhuis et al., 2021; Tisdall et al., 2022) and rodent models of tauopathies (Maphis et al., 2015; Sasaki et al., 2008). While the precise disease pathogenesis is not yet well understood, studies in animal models of tauopathy indicate that microgliosis may precede the overt formation of tau neurofibrillary tangles in the brain (Yoshiyama et al., 2007). Moreover, it is known that these changes at the subcellular level occur much before the onset of global tissue atrophy or cognitive deficits. Quantitative imaging markers to distinguish these alterations at the microscopic level therefore have substantive importance to assess pathological severity in specific brain regions and to understand the disease pathogenesis.

Quantitative susceptibility mapping (QSM) is an MRI technique sensitive to molecular changes in brain tissue that affect its local magnetic susceptibility (Liu et al., 2015; Schweser et al., 2016; Wang and Liu, 2014). Proteins such as tau tend to be diamagnetic due to the high concentration of paired electrons (Babaei et al., 2017; Gong et al., 2019). On the other hand, microglial cells express ferritin, the main iron-storage protein in the brain, which is paramagnetic (Langkammer et al., 2012). It has been shown that iron loading is a prominent feature of activated microglia in AD (Kenkhuis et al., 2021). In previous QSM studies, the detectability of tau neurofibrillary tangles in the brain has largely been attributed to increased iron content accompanying the tau deposits (Cogswell et al., 2021; Spotorno et al., 2020). A study in a mouse model of tauopathy similarly reported increased paramagnetic susceptibility in brain regions with low and intermediate tau pathology, whereas no significant susceptibility effects were observed in cortical areas with high tau burden (O’Callaghan et al., 2017). In a subsequent study using phantom experiments, Gong et al. (Gong et al., 2019) showed that tau protein has diamagnetic susceptibility. The same study also showed histology-like susceptibility contrasts reflecting iron and β-amyloid deposits in an AD mouse model. However, the differential effects of tau deposition and neuroinflammation or iron overload in the brain have not been reported.

QSM provides a measure of bulk magnetic susceptibility that represents the composite effect of molecular sources in an imaging voxel. However, in brain regions where sources of paramagnetic and diamagnetic susceptibility (e.g., iron and protein aggregates or myelin) co-localize in the same voxel, their opposing frequency contributions to the MR signal phase cannot be distinguished and may lead to underestimated susceptibility values. Therefore, conventional QSM methods cannot differentially quantify molecular sources with opposing susceptibilities in the same voxel (Schweser et al., 2011). More recently, different biophysical models have been proposed to disentangle sub-voxel contributions from paramagnetic and diamagnetic susceptibility sources (Chen et al., 2021; Dimov et al., 2022b; Li et al., 2023; Shin et al., 2021). These models consider a linear dependence of the reversible transverse relaxation rate on absolute magnetic susceptibility, which holds under the static dephasing regime approximation (Yablonskiy and Haacke, 1994). Such susceptibility source separation methods have shown promising applications to characterize multiple sclerosis lesions (Dimov et al., 2022a; Emmerich et al., 2021; Kim et al., 2023; Straub et al., 2023), and to investigate protein aggregation or demyelination in AD (Ahmed et al., 2023). The DECOMPOSE-QSM method proposed by Chen et al. (2021) uses three-pool complex exponentials to model the signal evolution based solely on multi-echo gradient echo (GRE) data and derive maps of paramagnetic component and diamagnetic component susceptibility. Disentangling the susceptibility source contributions at the sub-voxel level may potentially provide more specific quantification of tissue magnetic properties and their alterations due to pathological conditions.

In this study, we used paramagnetic and diamagnetic susceptibility source separation to investigate microstructural alterations in the brain in an established mouse model of tauopathy. The PS19-P301S tauopathy mouse model recapitulates the prominent features of human tauopathy, including gliosis, tau neurofibrillary tangle formation, neurodegeneration, and brain atrophy (Yoshiyama et al., 2007). We acquired multi-echo GRE data from fixed brains of adult PS19 transgenic and age-matched wild-type control mice at 11.7 T. The multi-echo data for the mouse brains were fit using a three-pool complex signal model (Chen et al., 2021), and parametric maps of paramagnetic and diamagnetic component susceptibility were derived. Group-wise analysis of the susceptibility maps was used to investigate regional differences between the tau transgenic and control mouse brains. The QSM and source separation maps were further compared with immunohistological staining for microglia and phosphorylated-tau, to decipher the underlying neuropathological changes contributing to the susceptibility differences.

## 2. METHODS AND MATERIALS

### 2.1 Animals and specimen preparation

All animal procedures were approved by the Animal Care and Use Committee at the Johns Hopkins University School of Medicine. Male and female transgenic PS19 mice (strain #008169) and age-matched wild-type (WT) control mice were obtained from the Jackson Laboratory (Bar Harbor, ME, USA). The PS19 transgenic tau mice (abbreviated as Tau mice) express the P301S mutant form of human tau protein driven by the mouse prion protein promoter, and develop age-related accumulation of tau neurofibrillary tangles. Tangle pathology is accompanied by microgliosis and neuronal loss, but not amyloid plaques (Yoshiyama et al., 2007). At 9 months of age, mice (n = 5 in each group) were transcardially perfused with phosphate buffered saline (PBS) and 4% paraformaldehyde (PFA) in PBS. After perfusion fixation, the mouse heads were placed in 4% PFA until imaging.

### 2.2 MRI acquisition

The specimens were transferred to PBS for five days before MRI, and placed in 20-mm diameter tubes that were filled with Fomblin (Solvay Inc., Princeton, NJ, USA), a proton-free perfluoropolyether, to prevent dehydration and for susceptibility matching at the tissue-air interface. MRI experiments were performed on an 11.7 T vertical-bore scanner (Bruker Biospin, Billerica, MA, USA) equipped with an actively-shielded Micro2.5 gradient system with a maximum gradient strength of 1500 mT/m. A 20-mm-diameter birdcage volume coil was used as the radiofrequency transceiver. The mouse brains were scanned with the dorsal-ventral axis of the skull oriented along the main magnetic field direction. A 3D multi-echo gradient echo (MGE) sequence with fast flyback for monopolar readout gradients was used for image acquisition. The imaging parameters were as follows: flip angle = 30°, pulse repetition time (TR) = 100 ms, first echo time (TE_1_) = 4 ms, 8 echoes with inter-echo spacing = 3.8 ms, 4 signal averages, and receiver bandwidth = 70 kHz. The 3D multi-echo data were acquired at an isotropic spatial resolution of 70 µm with a scan time of 3 h 45 min. The imaging field-of-view and acquisition matrix size were 12 mm x 9.2 mm x 17.6 mm and 170 x 132 x 252, respectively.

### 2.3 Image reconstruction and processing

The raw *k*-space data were reconstructed using custom-written scripts in IDL (ITT Visual Information Solutions, Boulder, CO) with zero-padding to twice the matrix size before Fourier transformation, and separated into magnitude and phase images (Aggarwal et al., 2018). The sum of squares of the GRE magnitude images from echoes 2 to 8 (TE = 7.8 to 30.6 ms) was used to extract the whole brain mask based on seeded region growing segmentation in RoiEditor (www.mristudio.org). The raw phase maps were unwrapped using Laplacian-based phase unwrapping, and the background phase was removed using the variable-kernel sophisticated harmonic artifact reduction for phase data (V-SHARP) method (Li et al., 2011) with a maximum spherical kernel radius of 25 voxels. Dipole inversion to calculate quantitative susceptibility maps was performed using the STAR-QSM method in StiSuite (Li et al., 2013; Wei et al., 2015). Quantitative susceptibility maps were also calculated by combining the multi-echo data for each mouse brain. The normalized phase φ was calculated from the unwrapped phase maps φ_*i*_, as (Wei et al., 2015):

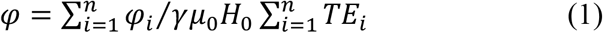

where *γ, μ*_0_, and *H*_0_ are the gyromagnetic ratio, vacuum permeability, and applied magnetic field, respectively, and *n* is the number of gradient echoes. The normalized background field was removed by V-SHARP filtering, followed by STAR-QSM to calculate the susceptibility maps. The susceptibility values in the QSM maps were intrinsically referenced to the mean susceptibility of the whole brain.

### 2.4 Model fitting for susceptibility source separation

To estimate sub-voxel contributions from paramagnetic and diamagnetic susceptibility sources, we used the DECOMPOSE-QSM method proposed by Chen et al. (2021). Briefly, the TE-dependent local complex signal at each voxel was synthesized as,

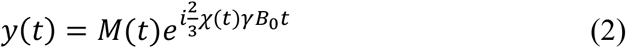

where *M*(*t*) is the magnitude of the raw signal and *χ*(*t*) is the quantitative susceptibility at each echo. *y*(*t*) was fit to a 3-pool model consisting of sub-voxel paramagnetic, diamagnetic, and neutral (*χ*_0_ = 0) susceptibility components. Under the static dephasing regime, the magnitude decay kernel representing the proportionality constant between the transverse relaxation rate 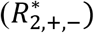 and volume susceptibility of paramagnetic or diamagnetic components (*χ*_+,−_) can be written as,

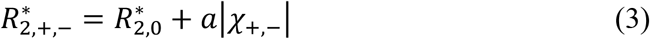

where 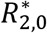 is the transverse relaxation rate of the reference susceptibility medium, and 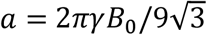 for spherical susceptibility sources (Chen et al., 2021; Yablonskiy and Haacke, 1994).

The theoretical value of *a* was calculated to be 1.26 kHz/ppm at 11.7 T. The total complex signal *S*(*t*) in the voxel with the three components can be written as:

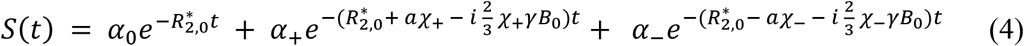

where *α*_0_, *α*_+_, and *α*_−_ denote the relative signal contributions of the reference, paramagnetic, and diamagnetic components at TE = 0 ms, respectively.

To estimate the linear and non-linear parameters in Eq. (4), the optimization problem was formulated as an alternating minimization problem with 3 steps similar to Chen et al. (2021):

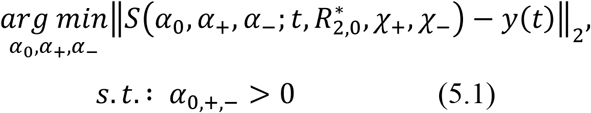

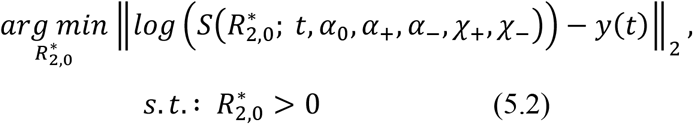

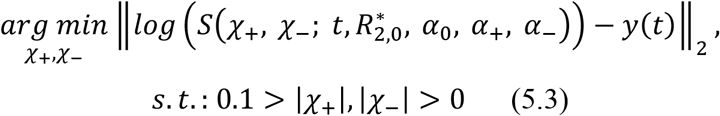

where ‖. ‖_2_ denotes the *L*2-norm. The solver was implemented in MATLAB (R2020a, Mathworks Inc., Natick, MA) using the *lsqcurvefit* function to solve for Eqs. (5.1) to (5.3) with manually calculated sparse block-diagonal Jacobians of the objective functions. 10 alternating iterations among the 3 steps were used to estimate the voxel-wise linear and non-linear parameters. Normalized signal fraction (*C*_0_, *C*_+_, *C*_−_) maps for the mouse brains were calculated from the fitted linear coefficients (*α*_0_, *α*_+_, *α*_−_) normalized to sum to one:

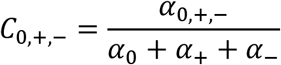

Composite paramagnetic component susceptibility (PCS) and diamagnetic component susceptibility (DCS) maps of the mouse brains were computed from the fitted model parameters according to Eqs. (6)-(8) of Chen et al. (2021).

### 2.5 Generation of group-specific templates

For quantitative analysis, group-averaged templates of the control WT and Tau mouse brains were generated. Briefly, the brain with the median volume in each group was chosen as the initial reference. The other brains in the group were aligned to the reference brain using rigid registration based on the magnitude images (sum of squares of the magnitude images from echoes 2 to 8), followed by 12-parameter linear affine registration (Woods et al., 1998). The registered images were intensity-averaged to generate the new reference. This process of affine registration and intensity averaging was repeated iteratively 5 times, with the reference image updated after each iteration and used as the target for subsequent registration (Aggarwal et al., 2009). The images in each group were then warped to the final reference using large deformation diffeomorphic metric mapping (LDDMM) (Beg et al., 2005) and averaged to generate the group-specific templates.

For structural annotation, the Allen mouse brain common coordinate framework (CCF) reference atlas (Wang et al., 2020) was non-linearly registered to the average group templates. The cross-modality registration was driven by similarity of gray matter-white matter contrasts between the GRE magnitude images and the CCF atlas, which is based on two-photon autofluorescence images of the mouse brain. The Allen atlas was first resampled to match the resolution of the MRI group templates, followed by affine registration and intensity histogram matching. Since the PS19 mice exhibit hippocampal atrophy, we used 2-channel LDDMM driven by magnitude images and a manually segmented binary mask of the hippocampus for improved registration accuracy. The combined diffeomorphic transformation, 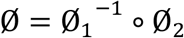, where Ø_1_ is the transformation from the atlas to the Tau group template, and Ø_2_ from the atlas to the control WT group template, was used to co-register all brains to the control group template space. The resulting transformations were then applied to warp the QSM and susceptibility source separation parametric maps of each subject to the common template space.

### 2.6 Statistical analysis

Cortical and subcortical regions-of-interest (ROIs) from the Allen CCF atlas were warped to the group-averaged templates and to the individual mouse brain images using the derived transformation matrices. Residual errors in the segmentation propagation were assessed visually and manually corrected based on the GRE magnitude images. Statistical comparisons of susceptibility values in the ROIs between the Tau transgenic and WT control mouse brains were conducted using Mann-Whitney U tests. All statistical calculations were performed using MATLAB, with the significance levels for the tests set as follows: ^**^*p*<0.01, and ^*^*p*<0.05. Data in the figures are presented as mean ± standard deviation (s.d.).

### 2.7 Immunohistochemistry

After MRI, the mouse brains were extracted from the skulls and processed for immunostaining. Fixed and sucrose-cryoprotected brains were sectioned coronally with a cryostat into 40-µm thick sections. Staining was performed on free-floating sections. The sections were rinsed 3 times for 10 min in PBS and were then incubated with blocking buffer (5% goat or donkey serum and 0.2% Triton X-100 in PBS) for 1 h at room temperature. Then the sections were incubated with primary antibodies in blocking buffer overnight at 4°C: AT8 (Mouse, 1:100, Thermo Scientific), and Iba1 (Rabbit, 1:1000, Wako; rat 1:100, Abcam). After thorough washes in PBS, sections were incubated with 1:1000 dilution of Alexa 488-, Alexa 555-, and Alexa 647-conjugated secondary antibodies (Thermo Scientific) appropriate for the species of the primary antibodies at room temperature for 1 h. The sections were examined and images acquired using a Zeiss LSM700 laser-scanning confocal microscope.

## 3. RESULTS

### 3.1 Susceptibility source separation in the mouse brain

Fig. 1 shows representative axial slices from the group-averaged QSM template of the WT mouse brains. The QSM template provided distinct contrasts highlighting iron-rich gray matter regions and neuronal layers, including the globus pallidus and layers in the hippocampus and cerebellum (Fig. 1a-d). The laminar structure of the hippocampus delineated based on susceptibility variations is clearly seen in Fig. 1. Fig. 1e further depicts markedly high paramagnetic susceptibility (13.2 ± 3.1 ppb) in the pyramidal cell layer of the hippocampal CA1 region, reflecting its high iron content. This layer is most distinctly highlighted in the medial CA1 in Fig. 1a-b, and appears relatively less paramagnetic in lateral hippocampal regions (Fig. 1c-d). Notably, this susceptibility pattern is consistent with a lateral-to-medial gradient of increasing iron concentration reported in the mouse CA1 pyramidal layer (Hackett et al., 2018).

**Fig. 1:**
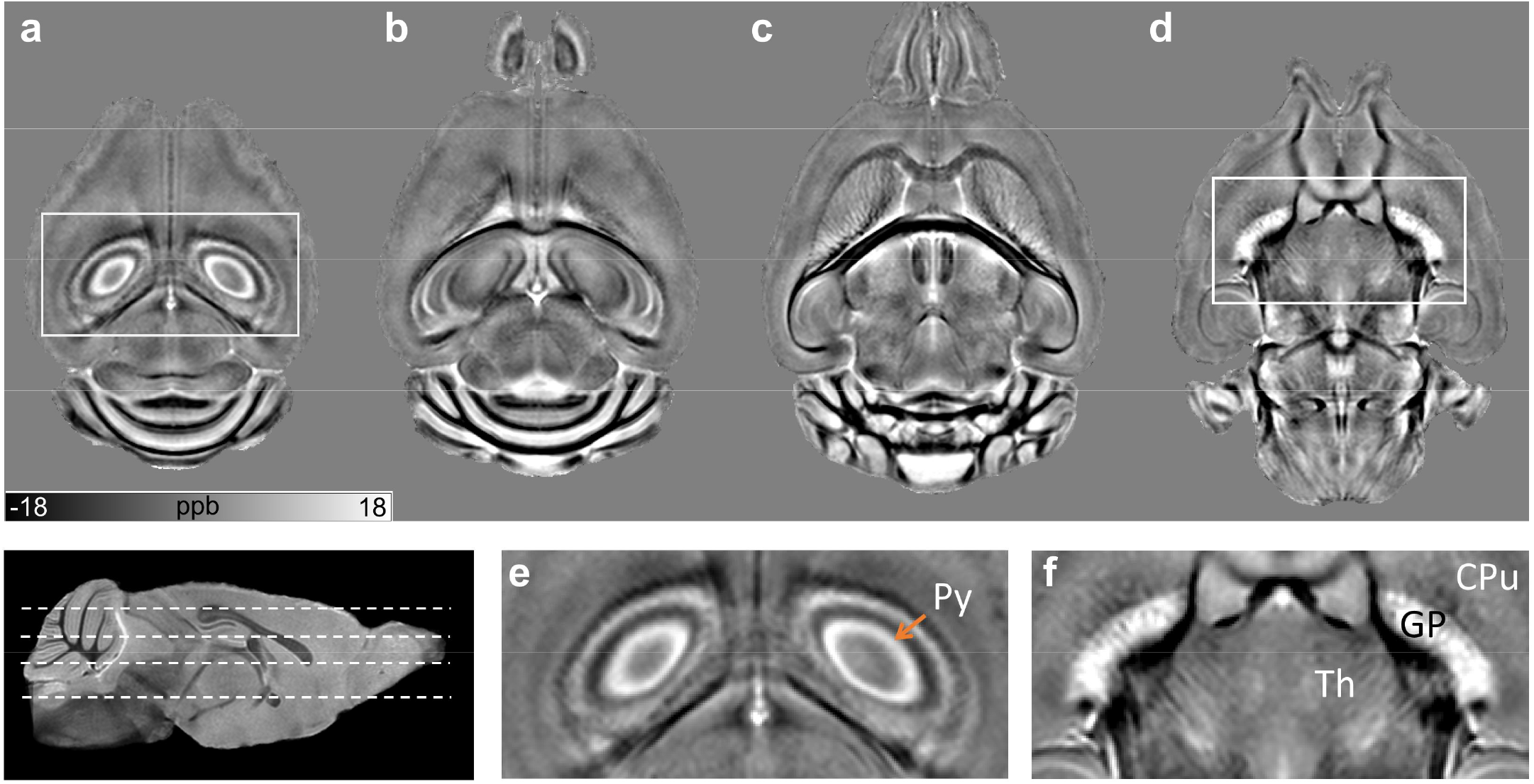
Group-averaged quantitative susceptibility template of the wild-type (WT) mouse brains. **a-d)** Representative axial slices from the QSM template of the WT group are shown, the locations of the slices are indicated by dashed lines in the scout magnitude image on the bottom left. **e, f)** Zoomed-in views of select regions (indicated by the white boxes) show distinctly highlighted neuronal layers and gray matter regions, including the hippocampal pyramidal cell layer (Py) and globus pallidus (GP), reflecting their high iron content. The laminar structure of the hippocampus delineated based on susceptibility variations is clearly seen in (e). Abbreviations: Th: thalamus, CPu: caudate putamen.

The resulting signal fraction (*C*_*0*_, *C*_*+*_, *C*_*-*_) and susceptibility source separation maps from fitting the three-pool signal model for the WT mouse brains are shown in Fig. 2. The *C*_*0*_, *C*_*+*_, and *C*_*-*_ maps represent the estimated fractional contributions of each of the three components to the signal at TE = 0 ms. The *C*_*0*_ map, which represents the neutral component fraction, appears relatively homogeneous in gray matter regions of the cortex and hippocampus (Fig. 2, top row). In comparison, the fitted paramagnetic (*C*_*+*_, PCS) and diamagnetic (*C*_*-*_, DCS) maps captured fine details of intra-cortical layers and neuronal layers in the hippocampus based on the intrinsic variations in susceptibility (zoomed-in views in Fig. 2). The diamagnetic signal fraction is seen to be high in white matter and myelinated brain regions, whereas the paramagnetic maps exhibit hyperintensity in iron-rich gray matter structures. The group-averaged PCS and DCS maps in Fig. 2 (bottom row) exhibit distinct anatomical contrasts that predominantly reflect known variations in iron and myelin content in the mouse brain, respectively.

**Fig. 2:**
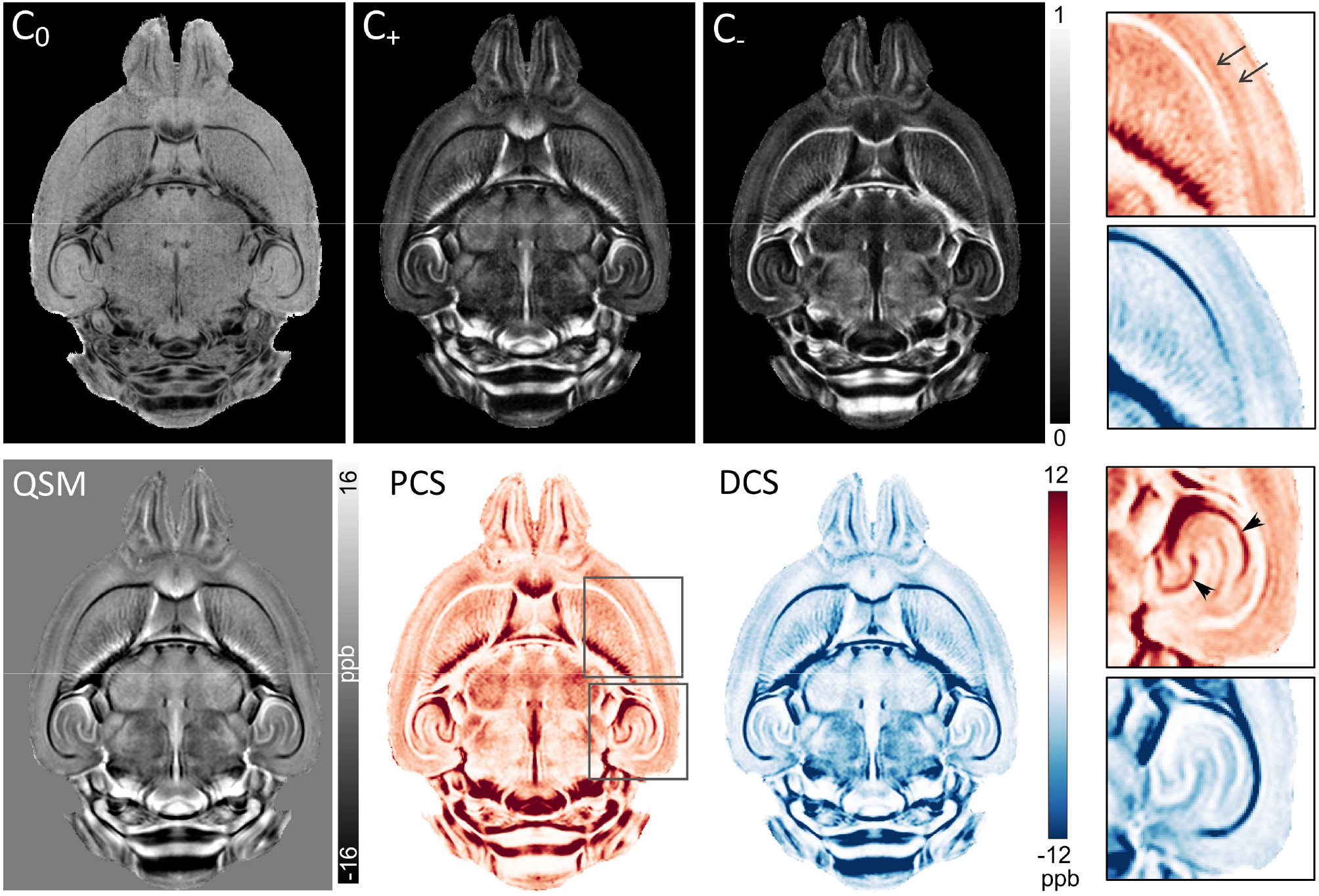
Parametric maps from fitting the three-pool complex signal model for QSM source separation (Eq. 4) to data for the WT mouse brains. A representative mid-axial slice is shown from the group-averaged signal fraction (C_0_, C_+_, C_-_) maps (top row), and the QSM, paramagnetic component susceptibility (PCS), and diamagnetic component susceptibility (DCS) maps (bottom row). The PCS and DCS maps exhibit distinct anatomical contrasts reflecting known variations in iron and myelin content in the mouse brain, respectively. Zoomed-in views of select regions indicated by the boxes (right panel) reveal layers in the cerebral cortex (arrows) and the granule and pyramidal cell layers in the hippocampus (arrowheads) distinguished in the PCS map.

### 3.2 Regional variations in paramagnetic and diamagnetic susceptibility in Tau mice

Fig. 3 compares the QSM and fitted DECOMPOSE model parameter maps for the WT and Tau transgenic mouse brains at the level of the dorsal hippocampus. Marked differences between the parametric maps of the two groups are seen in the hippocampus. The WT maps showed a distinct laminar structure in the hippocampus, with high paramagnetic susceptibility in the pyramidal cell layer and dentate gyrus, and high diamagnetic susceptibility in the stratum lacunosum moleculare (slm) that contains myelinated fibers. In comparison, maps of the PS19 Tau mice revealed diffusively elevated paramagnetic susceptibility in the dorsal hippocampus, with a disruption of this laminar structure (Fig. 3a-b). The localized PCS increase in the hippocampus of the Tau mice is clearly seen in Fig. 3 (Fig. 3b, arrowheads). In addition, the DCS maps depict lower diamagnetic susceptibility in the slm of the Tau mice relative to WT mice, likely reflecting axonal degeneration in the hippocampus previously shown to occur in this model (Yoshiyama et al., 2007).

**Fig. 3:**
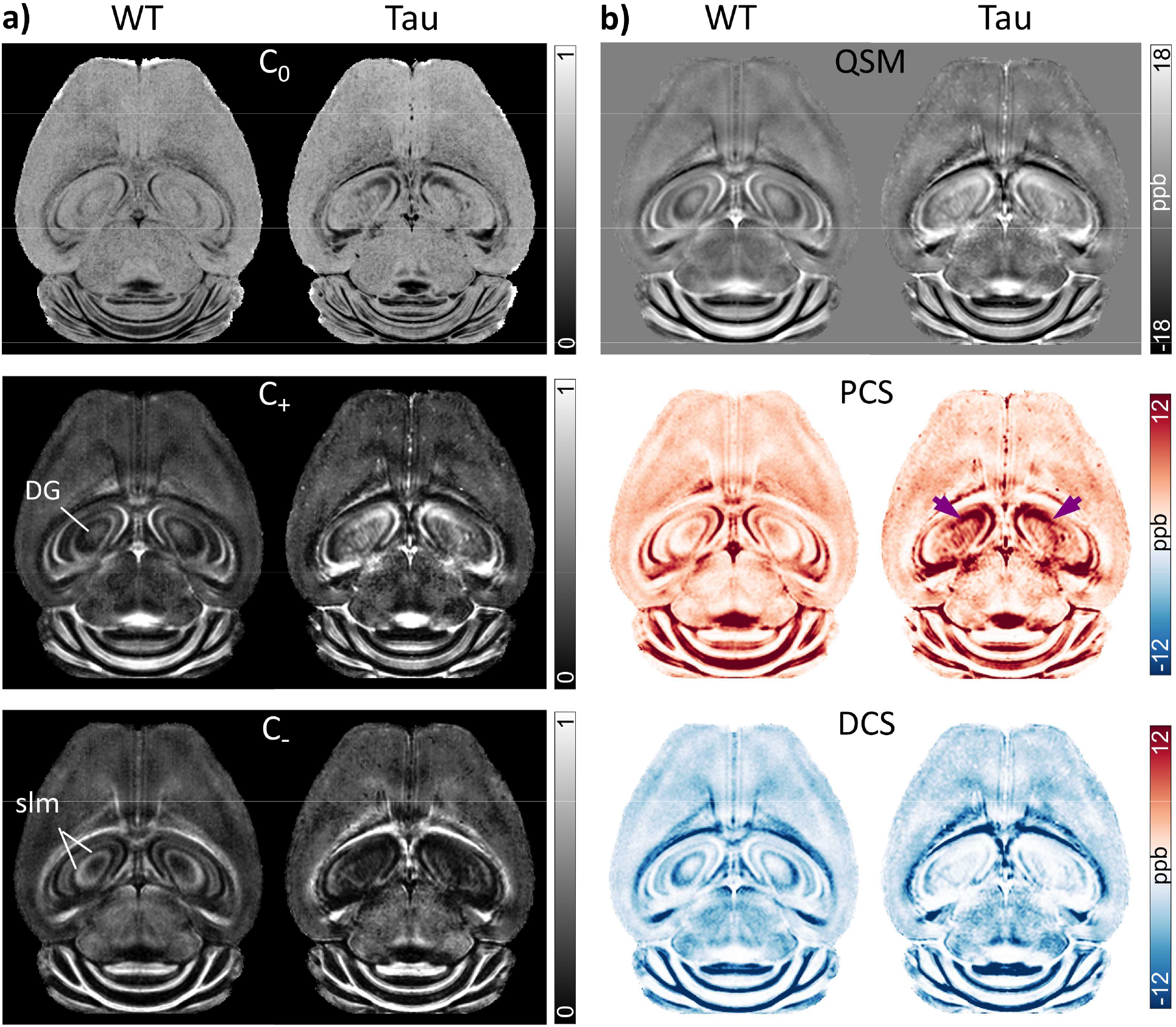
Comparison of group-averaged QSM and susceptibility source separation maps of the WT and Tau mouse brains at the level of the dorsal hippocampus. A representative axial slice from (a) signal fraction (C_0_, C_+_, C_-_) maps and (b) QSM, PCS, and DCS maps in template space is shown. Maps of the WT group show a distinct laminar structure in the hippocampus, with high paramagnetic susceptibility in the Py and dentate gyrus (DG), and high diamagnetic susceptibility in the stratum lacunosum moleculare (slm). In comparison, maps of the PS19 group reveal diffusely elevated paramagnetic susceptibility in the dorsal hippocampus (arrowheads in the PCS map).

Quantitative PCS values for different sub-regions of the dorsal hippocampus are plotted in Fig. 4. Significantly higher mean PCS values were observed in the dorsal subiculum (*p* = 0.0079), dentate gyrus (*p* = 0.0079), and CA1 (*p* = 0.016) regions of the Tau mice relative to WT controls. In comparison, no significant differences between the two groups were observed in the dorsal somatosensory and motor cortical areas (Fig. 4). Diffusely elevated PCS was observed in the Tau mouse brains throughout the dorsal-ventral extent of the hippocampus, especially in caudal regions. A mid-axial slice from the QSM maps of the two groups in shown in Fig. 5, together with zoomed-in views of the PCS and DCS maps. The maps exhibit bilaterally elevated paramagnetic susceptibility in caudal aspects of the PS19 hippocampus compared to corresponding regions in the WT brains (Fig. 5a, b).

**Fig. 4:**
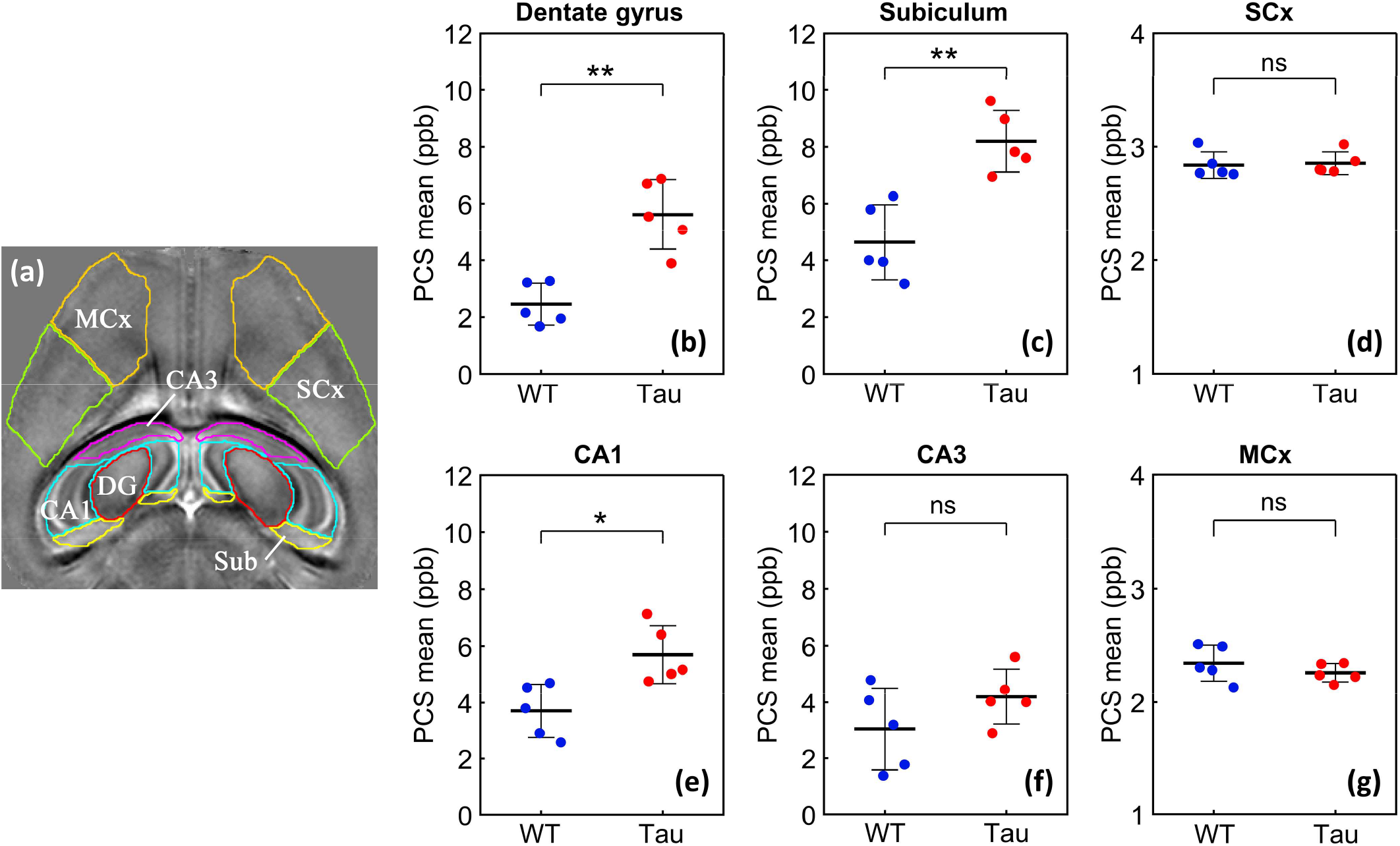
Mean PCS values in sub-regions of the dorsal hippocampus for the WT and Tau mouse brains. **a)** Hippocampal and cortical regions of interest (ROIs) from the Allen atlas are overlaid on a representative axial slice from the WT QSM template. DG: dentate gyrus, Sub: subiculum, MCx: primary motor cortex, SCx: somatosensory cortex. **b-g)** Plots of mean PCS values for the WT (blue circles) and Tau (red circles) mice. Data points denote the mean PCS values measured across five consecutive slices in the dorsal hippocampus of each mouse. Horizontal lines indicate the mean (± s.d.) for each group. Significance levels for group differences are indicated as; ^**^*p* = 0.0079, ^*^*p* = 0.016, ns: not significant (Mann-Whitney U test, n = 5). Note that the y-axis for the cortical ROI plots is stretched by a factor of 4 to better visualize the degree of inter-subject variability.

**Fig. 5:**
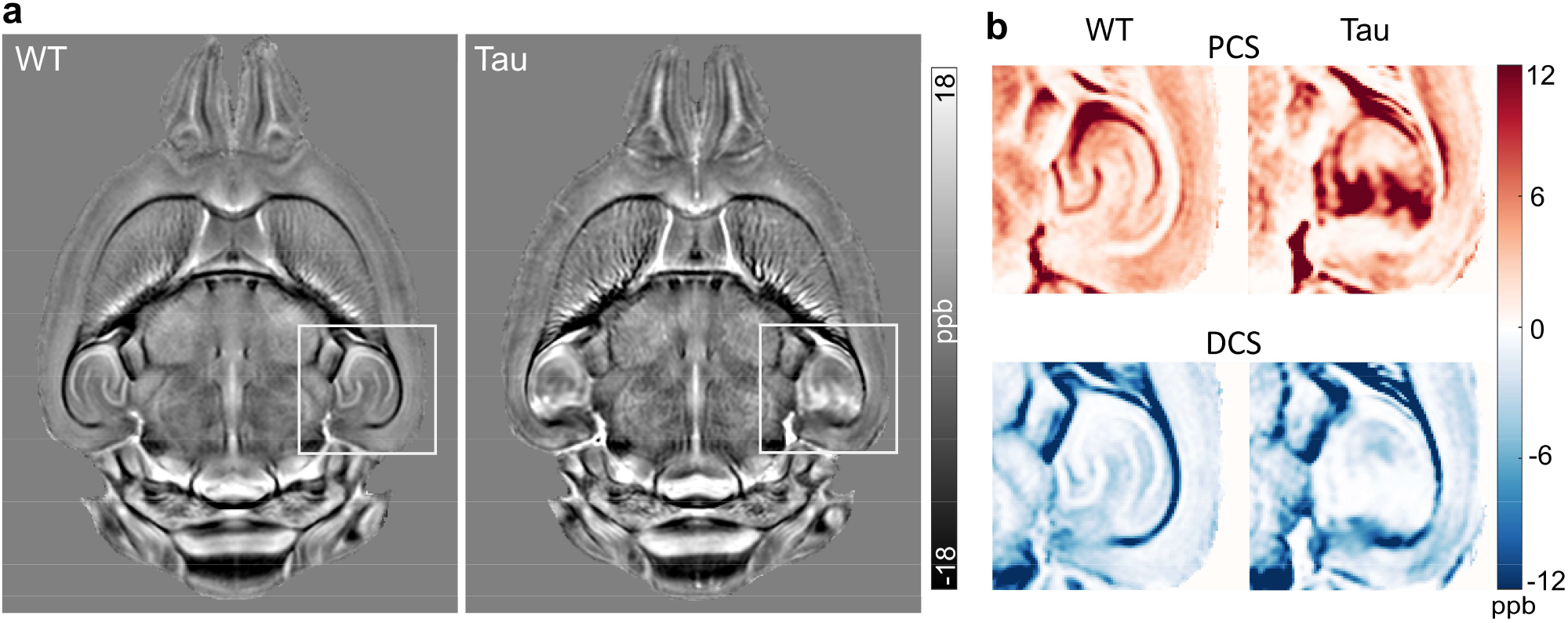
Comparison of QSM and susceptibility source separation maps of the WT and Tau mouse brains at the mid-axial level. **a)** A representative axial slice is shown from the group-averaged QSM maps in template space, revealing bilateral hyperintense areas with higher paramagnetic susceptibility in the hippocampus of the Tau mice. **b)** Zoomed-in views of PCS and DCS maps of the regions indicated by the white boxes in (a).

Fig. 6 compares an axial slice from the group-averaged QSM and DCS maps at the level of the entorhinal cortex. Focal increases in paramagnetic susceptibility in the PS19 hippocampus were also observed at this level. Additionally, the maps revealed bilateral areas with markedly higher diamagnetic susceptibility (hypointense areas in the QSM maps in Fig. 6a) in the entorhinal cortex of PS19 mice compared to corresponding regions in the WT group. The regionally elevated |DCS| in the entorhinal cortex of the Tau mice is clearly seen in Fig. 6 (Fig. 6b, arrows).

**Fig. 6:**
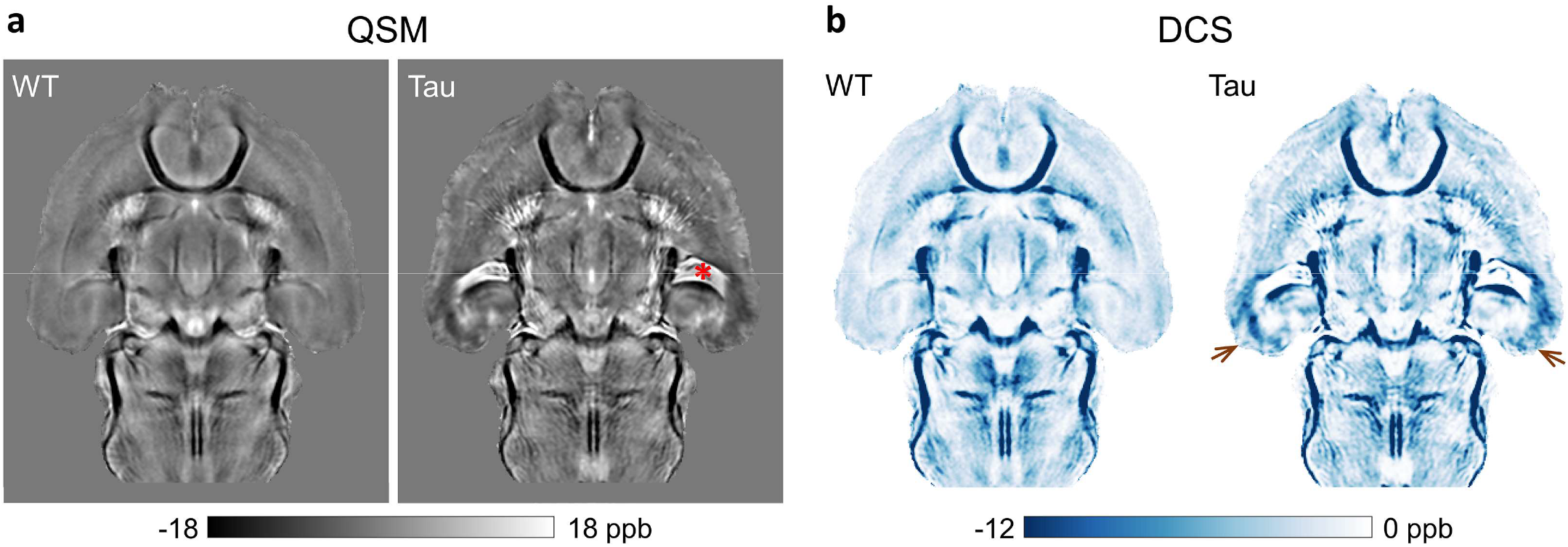
Comparison of group-averaged QSM and DCS maps of the WT and Tau mouse brains at the level of the entorhinal cortex. **a)** An axial slice from the QSM maps in template space, revealing focal increases in paramagnetic susceptibility (hyperintense areas) and diamagnetic susceptibility (hypointense areas) in the hippocampus and entorhinal cortex of the Tau mice, respectively. Enlarged ventricles are also seen in the Tau group (asterisk), which appear paramagnetic likely due to high iron content in the choroid plexus. **b)** DCS maps of the two groups, showing bilaterally elevated |DCS| in the entorhinal cortex of the Tau mice (arrows) compared to the WT mice.

Representative coronal slices from the group-averaged GRE magnitude, QSM, PCS, and DCS maps of the WT and Tau mouse brains are shown in Figs 7-8. Fig. 7 exhibits bilaterally elevated paramagnetic susceptibility in the caudal hippocampus of the Tau mice compared to the WT mice, in regions corresponding to the subiculum, CA1, and dentate gyrus. The PCS increase is consistent with extensive microgliosis previously reported in the hippocampus of the PS19 mouse brains (Yoshiyama et al., 2007). The PCS difference map between the two groups (Fig. 7d) further shows the regional differences localized to the hippocampus, and to a lesser extent, the piriform/entorhinal cortex, as compared to adjacent gray matter regions. Fig. 8 shows a more caudal slice, depicting markedly higher diamagnetic susceptibility in the entorhinal cortex, presubiculum, and subiculum areas of the PS19 mice compared to WT mice. The region-specific differences between the two groups can be clearly distinguished in the DCS maps (Fig. 8c, d).

**Fig. 7:**
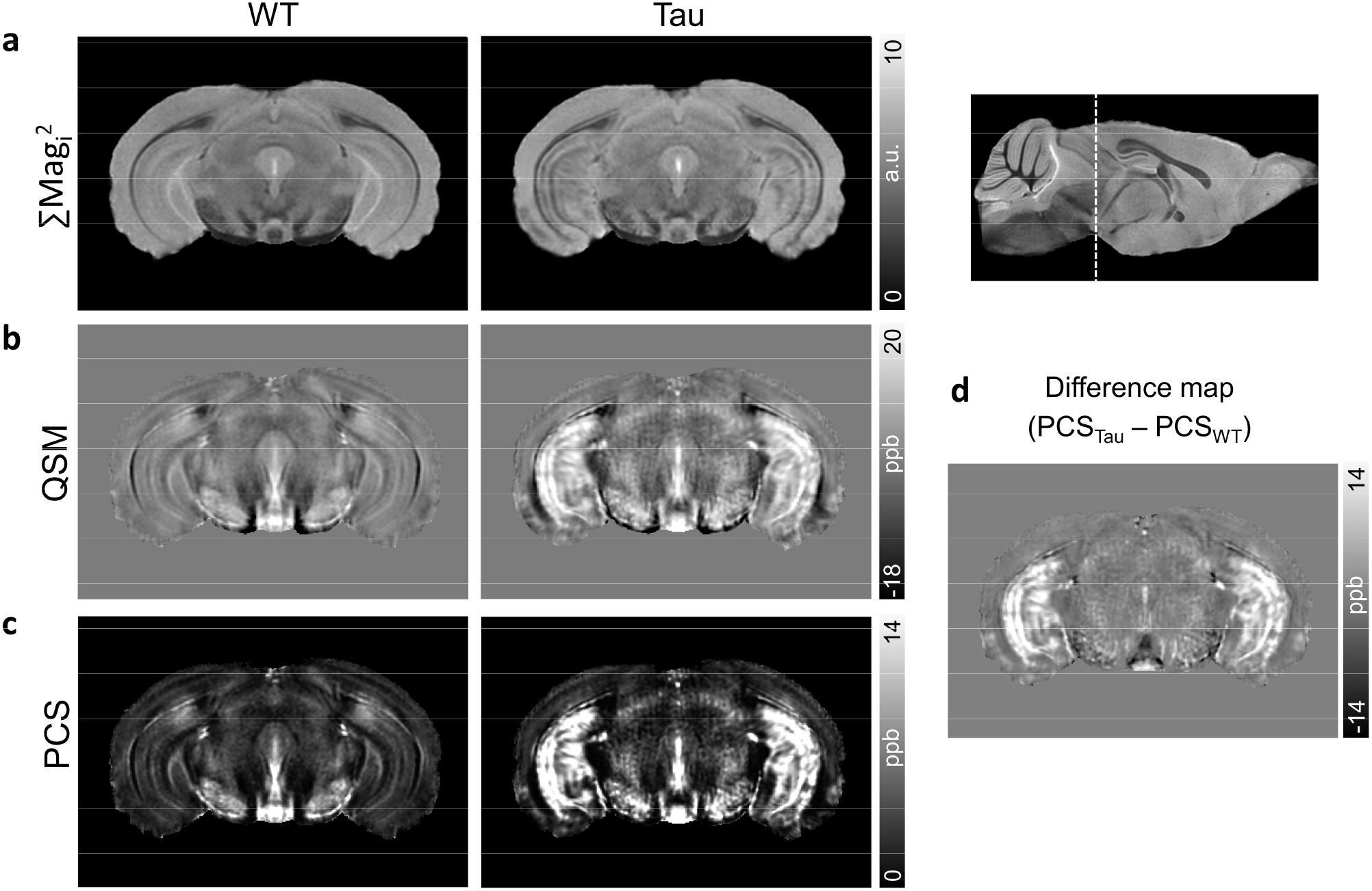
A coronal slice from the group-averaged GRE magnitude (a), QSM (b), and PCS maps (c) of the WT and Tau mouse brains in template space, at the level of the caudal hippocampus. The anatomical location of the slice is indicated by the dashed line in the sagittal scout image at the top right. The QSM and PCS maps reveal bilateral areas in the hippocampus of the Tau mice with markedly higher paramagnetic susceptibility relative to corresponding regions in the WT mice. The PCS difference map between the two groups (d) reveals pronounced PCS differences specifically in the hippocampus as compared to adjacent gray matter regions.

**Fig. 8:**
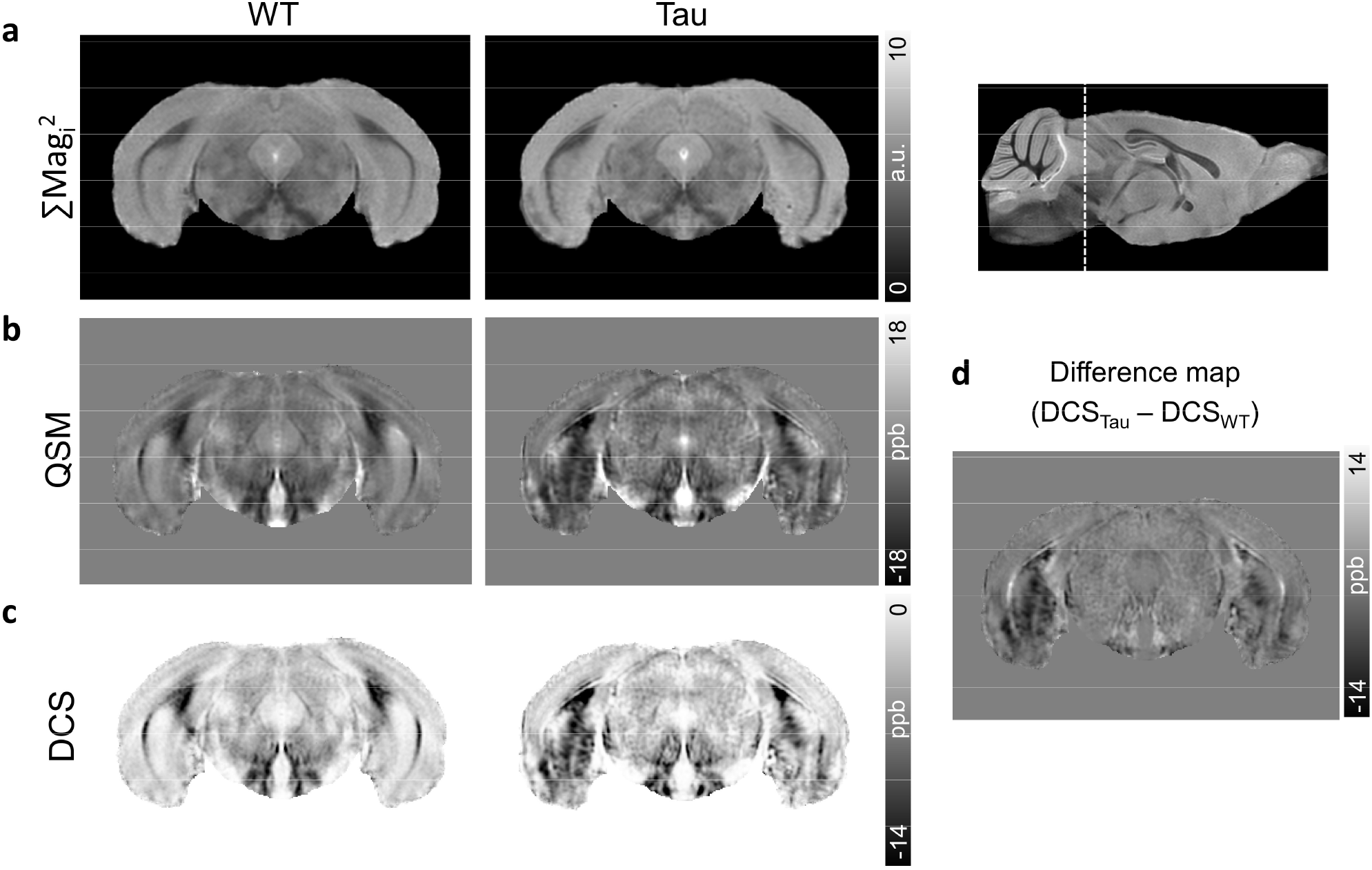
A coronal slice from the group-averaged GRE magnitude (a), QSM (b), and DCS maps (c) of the WT and Tau mouse brains in template space, taken at the level of the entorhinal cortex. The QSM and DCS maps reveal bilateral regions with high diamagnetic susceptibility (hypointense areas) in the entorhinal cortex, presubiculum, and subiculum of the Tau mice compared to corresponding regions in the WT control mice. The DCS difference map between the two groups (d) shows the DCS differences localized to specific areas of the entorhinal cortex and hippocampus as compared to adjacent gray matter regions.

### 3.3 ROI analysis

Fig. 9 shows structural delineations from the Allen mouse atlas overlaid on QSM maps of the WT and Tau brains, together with plots of DCS and PCS values in select regions. The mean PCS and DCS values for the ROIs averaged across five coronal slices at the level of the caudal hippocampus are listed in Table 1. Susceptibility values for the globus pallidus, an iron rich region, are also shown for comparison. The subiculum, CA1, CA3, and dentate gyrus sub-regions of the caudal hippocampus exhibited significantly higher PCS (*p* < 0.01) in the Tau mouse brains than corresponding regions in the WT mouse brains. In comparison, significantly higher |DCS| was observed in the entorhinal cortex, postpiriform transition area, and cortical amygdala of the Tau mice relative to corresponding regions in the WT mice (Table 1). Interestingly, the postpiriform transition area showed both significantly higher PCS (*p* < 0.01) and significantly higher |DCS| (*p* < 0.05) in the Tau mice relative to the WT mice (Table 1). Conversely, QSM values for the two groups did not differ significantly in the postpiriform transition area. The substantia nigra, which has high iron content and thus high PCS, showed no significant susceptibility differences between the two groups (Fig. 9 and Table 1).

**Table 1:**
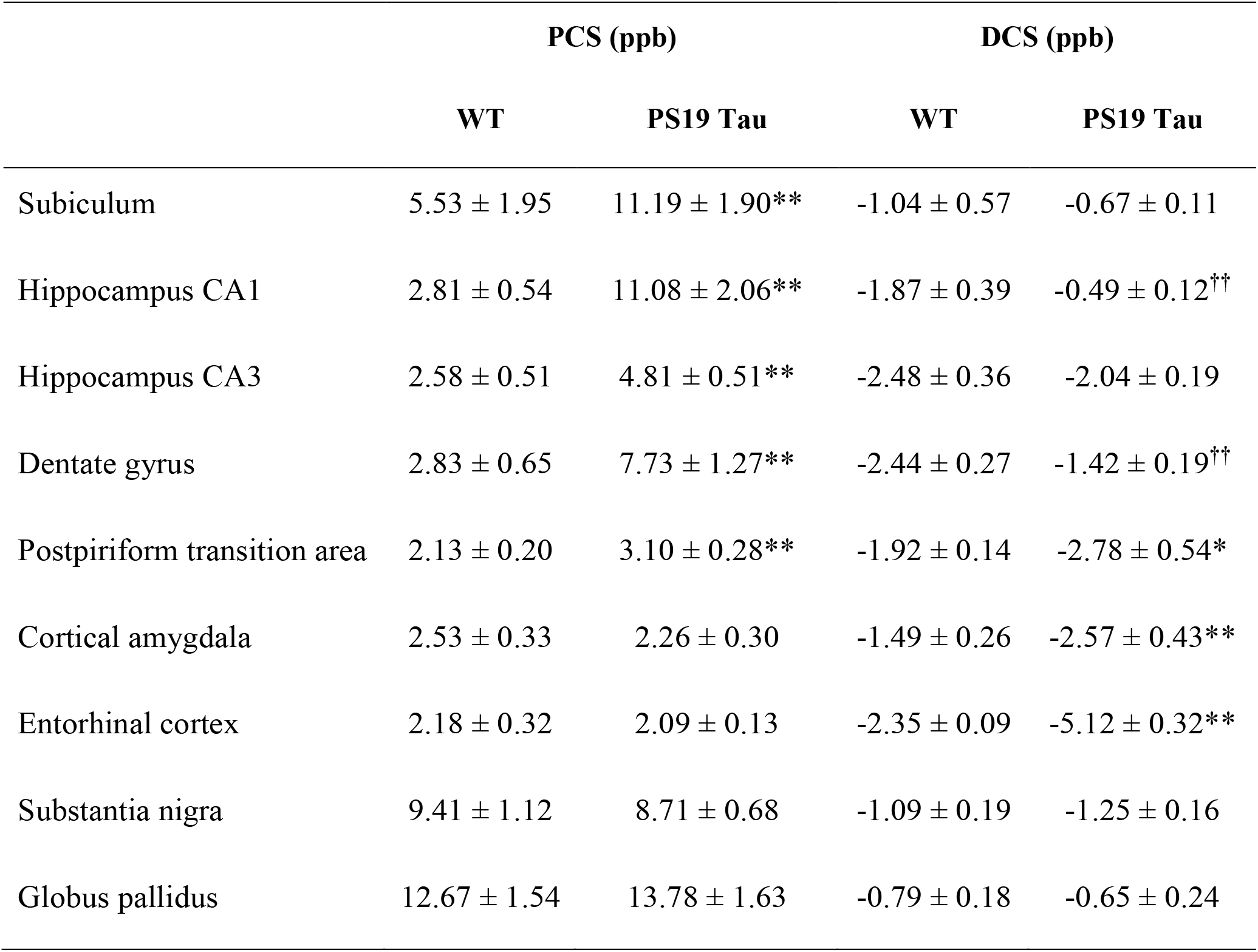
Mean (± s.d.) PCS and DCS values in gray matter ROIs in the WT and Tau mouse brains measured across five consecutive coronal slices. Significant group differences are denoted as: ^**^*p* < 0.01, ^*^*p* < 0.05 (Two-tailed Mann-Whitney U test, n = 5). ^††^ indicates significant (*p* < 0.01) group differences with lower regional |DCS| values in the Tau group than the WT group.

|  | PCS (ppb) |  | DCS (ppb) |  |
| --- | --- | --- | --- | --- |
|  | WT | PS19 Tau | WT | PS19 Tau |
| Subiculum | 5.53 ± 1.95 | 11.19 ± 1.90** | -1.04 ± 0.57 | -0.67 ± 0.11 |
| Hippocampus CA1 | 2.81 ± 0.54 | 11.08 ± 2.06** | -1.87 ± 0.39 | -0.49 ± 0.12<sup>††</sup> |
| Hippocampus CA3 | 2.58 ± 0.51 | 4.81 ± 0.51** | -2.48 ± 0.36 | -2.04 ± 0.19 |
| Dentate gyrus | 2.83 ± 0.65 | 7.73 ± 1.27** | -2.44 ± 0.27 | -1.42 ± 0.19<sup>††</sup> |
| Postpiriform transition area | 2.13 ± 0.20 | 3.10 ± 0.28** | -1.92 ± 0.14 | -2.78 ± 0.54* |
| Cortical amygdala | 2.53 ± 0.33 | 2.26 ± 0.30 | -1.49 ± 0.26 | -2.57 ± 0.43** |
| Entorhinal cortex | 2.18 ± 0.32 | 2.09 ± 0.13 | -2.35 ± 0.09 | -5.12 ± 0.32** |
| Substantia nigra | 9.41 ± 1.12 | 8.71 ± 0.68 | -1.09 ± 0.19 | -1.25 ± 0.16 |
| Globus pallidus | 12.67 ± 1.54 | 13.78 ± 1.63 | -0.79 ± 0.18 | -0.65 ± 0.24 |

**Fig. 9:**
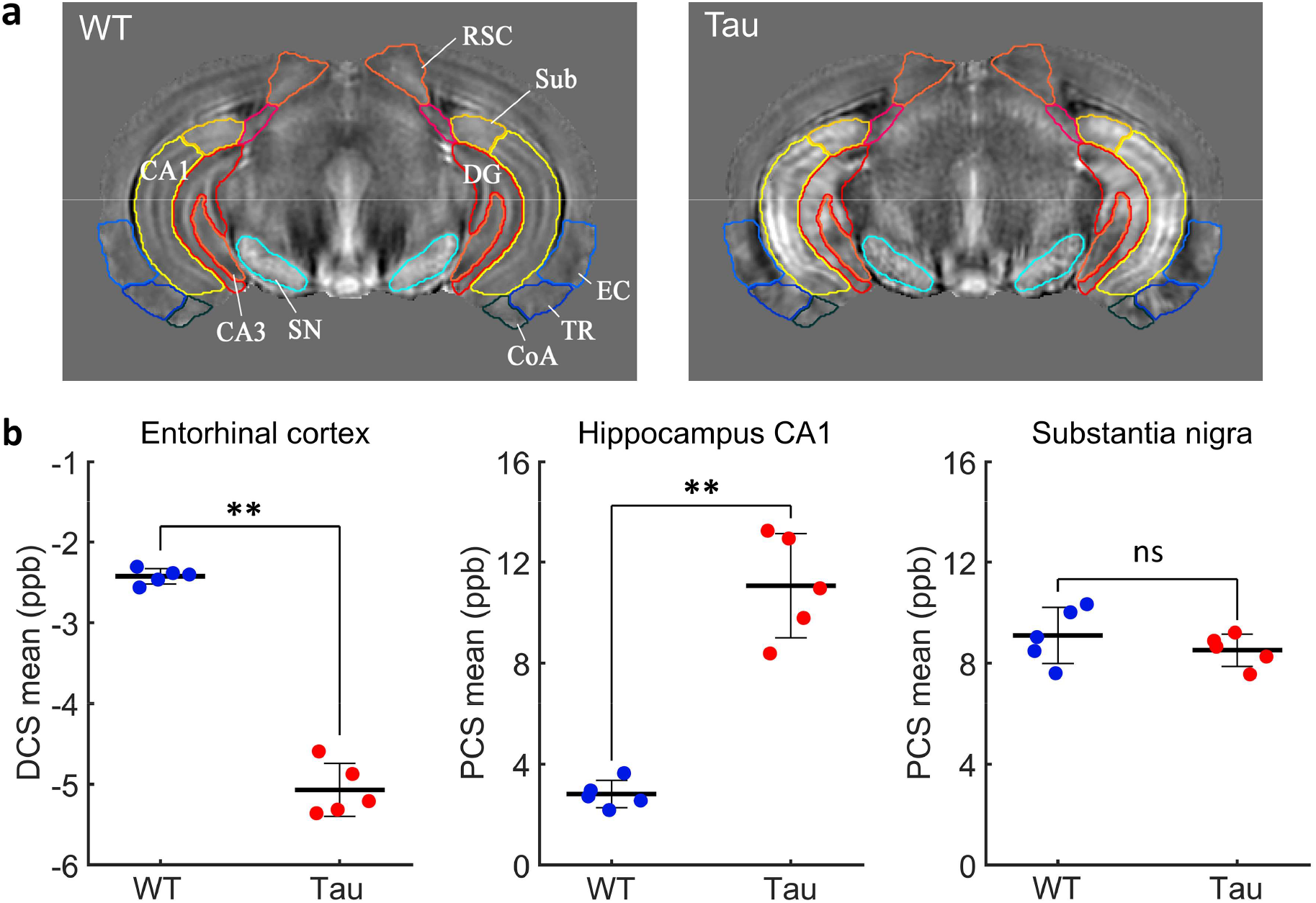
**a)** Structural delineations from the Allen atlas overlaid on the group-averaged QSM templates of the WT and Tau mouse brains at the level of the caudal hippocampus. **b)** Plots showing the mean DCS and PCS values in select ROIs for five mouse brains in each group. Data points represent the mean value for each mouse taken over five coronal slices at the level of the caudal hippocampus. Horizontal bars denote the mean (± s.d.) for each group. Significance levels for the group differences are indicated as; ^**^*p* = 0.0079, ^*^*p* = 0.016, ns: not significant (Mann-Whitney U test, n = 5). CoA: cortical amygdala, DG: dentate gyrus, EC: entorhinal cortex, RSC: retrosplenial cortex, SN: substantia nigra, Sub: subiculum, TR: postpiriform transition area.

### 3.4 Immunohistochemistry of WT and Tau brains

Fig. 10 shows coronal sections from representative WT and PS19 mouse brains immunostained for microglia (Iba1) and phosphorylated tau (AT8). The WT brains exhibited sparse Iba1-positive microglial cells with ramified non-reactive morphology in the hippocampus and cortex. In comparison, the PS19 brains exhibited a marked increase in the number of Iba1-positive microglia with swollen cell somas, indicative of extensive microgliosis in the subiculum, dentate gyrus, and CA1 regions of the hippocampus and in ventral cortical areas (Fig. 10). The PS19 mouse brains showed strong AT8-positive neuronal staining, indicative of phosphorylated tau deposits, in the cortical amygdala, entorhinal cortex, and postpiriform transition area (Fig. 10c), and relatively weaker staining in the hippocampal dentate gyrus and CA1 (Fig. 10b). No AT8-positive staining was observed in the WT mouse brains (Fig. 10b-c). The spatial extent of microgliosis in the hippocampus and dominant phosphorylated-tau deposits in the entorhinal cortical area in Fig. 10 correspond closely to the regional PCS and DCS differences between the Tau and WT mouse brains.

**Fig. 10:**
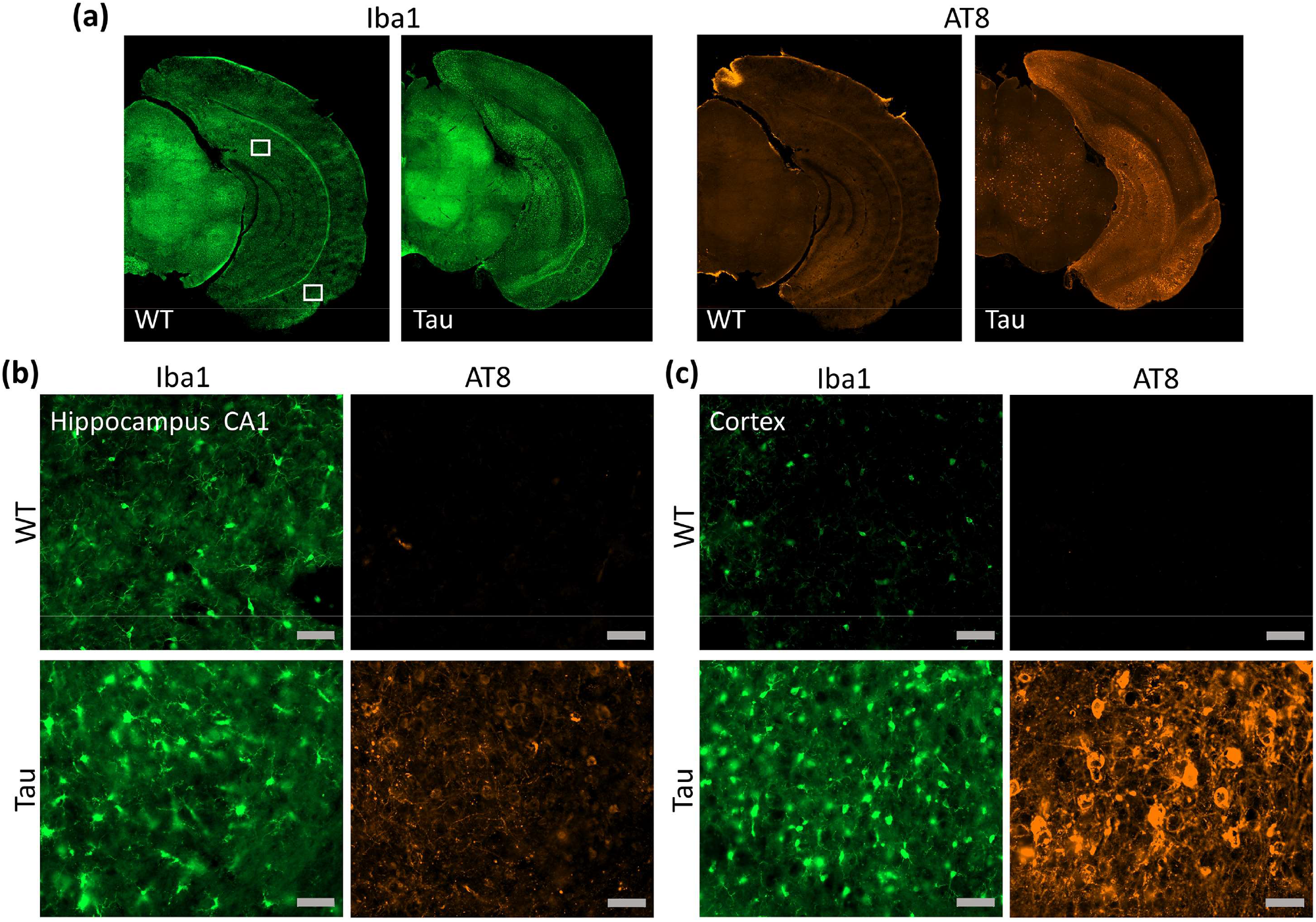
Sections from WT and PS19 Tau mouse brains stained for Iba1 (microglia) and AT8 (phosphorylated tau). (a) Coronal sections at the level of the caudal hippocampus from representative mouse brains in each group. (b-c) High-magnification sections of select regions in the hippocampus CA1 (b) and the entorhinal cortex/postpiriform transition area (c). The locations of the sections are indicated by boxed areas in (a). Scale bars in (b, c) = 50 µm.

## 4. DISCUSSION

This study demonstrates that QSM and susceptibility source separation can differentiate localized pathological alterations in the mouse brain with tauopathy. PCS and DCS maps provided distinct contrasts reflecting the known variations in iron and myelin content in the WT mouse brain, respectively. Group-wise analysis revealed significant regional differences between parametric maps of the WT and PS19 Tau mouse brains. Localized increases in absolute PCS and DCS were observed in specific hippocampal and cortical sub-regions of the PS19 mice relative to WT controls, which were found to correspond to regional microgliosis and tau deposition seen with immunohistology. The findings show the potential of quantitative susceptibility source separation to distinguish sub-cellular pathological alterations in the mouse brain. The ability to distinguish these pathological processes with MRI has key potential applications for mapping of disease severity and progression in specific brain regions and understanding the disease pathogenesis.

As seen in Figs. 1 and 2, QSM and source separation maps of the WT mouse brains showed a distinct laminar structure in the hippocampus, with delineation of iron-rich neuronal cell layers. The delineation of hippocampal layers in QSM maps was also shown in a previous study by Wei et al. (Wei et al., 2016). The pyramidal cell layer of the hippocampus contains densely-packed cell somas, that are shown to have high intracellular iron content (Sands et al., 2016). Interestingly, our study revealed a marked lateral-to-medial gradient of increasing paramagnetic susceptibility in the CA1 pyramidal cell layer of the WT mice. A previous study using X-ray fluorescence imaging of mouse brain sections also reported a lateral-to-medial gradient of increasing iron concentration in the CA1 pyramidal neurons, and suggested that the gradient of iron concentration may underlie the pattern of selective vulnerability of pyramidal neurons observed in many neurodegenerative diseases (Hackett et al., 2018). Mapping the 3D gradient of iron concentration with QSM may therefore enable further insights into the role of intracellular iron in the region-specific vulnerability of pyramidal neurons observed in neurodegenerative diseases.

Our study showed significant DCS differences between the WT and Tau mouse groups in regions including the cortical amygdala, entorhinal cortex, and piriform cortical areas, that are known to be severely affected by tau pathology (Hurtado et al., 2010; Ramirez et al., 2023). Previous studies have largely reported increased paramagnetic susceptibility associated with tau pathology, in both human brain and animal models. Studies in human AD patients showed positive correlations between regional susceptibility and tau positron emission tomography (tau-PET) standardized uptake value ratios in the basal ganglia and cortices (Cogswell et al., 2021; Spotorno et al., 2020), which were partly attributed to co-localization of iron or off-target binding of tau PET ligands. In the rTg4510 mouse model of tauopathy, O’Callaghan et al. (2017) showed higher paramagnetic susceptibility in brain regions with low tau deposits and neuroinflammation, while no significant susceptibility effects were observed in cortical areas with the highest tau burden. In a recent study, Ahmed et al. (Ahmed et al., 2023) showed negative associations between mean |DCS| and tau-PET signal in several limbic and cortical regions in AD. In the current study, significantly higher |DCS| was observed in the Tau mouse brains compared to the WT brains in the entorhinal cortex and parahippocampal regions. The regional differences in |DCS| corresponded closely to dominant AT8-positive staining for phosphorylated-tau deposits in these regions in the PS19 mice. These results indicate that tau deposits have a strong diamagnetic effect on tissue susceptibility. In regions were tau aggregates and inflammation or iron overload co-localize, separation of QSM into paramagnetic and diamagnetic components may provide improved specificity to distinguish these processes. In the postpiriform transition area, mean PCS and mean |DCS| were both significantly higher in the Tau mouse brains relative to the WT group. However, mean QSM values in this region did not differ significantly between the two groups.

In addition to tau deposits, DCS maps can also be affected by changes in myelin content. DCS contrast in the WT mouse brains is largely driven by myelin content (Liu et al., 2011). We also observed significantly lower |DCS| specifically in the CA1 and dentate gyrus of the Tau mice compared to controls (Table 1), which may reflect axonal degeneration or myelin loss in these regions. Reduction of mossy fibers has previously been reported in the dentate gyrus of PS19 mouse brains (Yoshiyama et al., 2007), and it was shown that synaptic pathology and axonal degeneration precede the formation of tau tangles in the hippocampus. Notably, myelin loss is expected to have an opposite effect on DCS than tau aggregation, and may be one possible explanation for the |DCS| decrease seen in the dentate gyrus and CA1 sub-regions. Further histological studies are, however, needed to disentangle the potential contribution of hippocampal myelin loss or axonal degeneration to the DCS maps.

Hippocampal and cortical regions with significantly higher PCS in the PS19 mice in our study showed a marked increase in Iba1-positive cells with swollen cell somas, indicative of microgliosis. Our findings are consistent with a previous study by O’Callaghan et al. (2017), that showed higher susceptibility in regions with increased staining for reactive microglia and astrocytes in a mouse model of tauopathy. MRI-histopathology studies in human AD and tauopathies have also reported association of T2^*^-weighted hypointensity with accumulation of iron-laden microglia (Tisdall et al., 2022; Zeineh et al., 2015). Iron loading has been shown to be a prominent feature of activated microglia in AD (Kenkhuis et al., 2021). Upregulation of the intracellular iron-storage protein ferritin light chain has been reported in AD, and studies have shown that the ferritin-positivity is specifically associated with microglia (Connor et al., 1992; Grundke-Iqbal et al., 1990), particularly microglia in dystrophic states (Lopes et al., 2008). Ferritin immunohistochemistry has also been suggested as a marker for dystrophic microglia (Shahidehpour et al., 2021). Although the precise mechanism is complex and not yet well understood, evidence indicates that the interaction between microgliosis and iron overload may be bidirectional (Long et al., 2022). The bilateral increase in PCS in the hippocampus of PS19 mice observed in our study, as seen in Fig. 7, is closely consistent with the spatial extent of microglial activation seen with autoradiography studies in this model (Maeda et al., 2011; Yoshiyama et al., 2007). These results suggest that PCS may provide a sensitive imaging marker for microglia-mediated inflammation and iron dyshomeostasis in the Tau mouse brains.

Regional iron accumulation can contribute to free radical generation, leading to oxidative stress and exacerbation of tau pathology (Derry et al., 2020). Previous studies have shown that microglial activation precedes the overt formation of tau tangles in the PS19 mouse brain, and microgliosis may contribute to the spread of tau pathology along anatomically connected brain regions (Maphis et al.,2015). The ability to track changes in PCS and DCS at different time points may therefore provide valuable insights into the disease pathogenesis.

It is also important to note some limitations of the study. QSM source separation is based on modeling a linear dependence of the transverse relaxation rate on absolute susceptibility in the static dephasing regime. The theory of static dephasing holds for sufficiently large magnetic susceptibility inclusions where the GRE signal dephasing is not affected by molecular diffusion. The validity of this regime depends on the field strength (*B*_0_), intrinsic diffusivity, and the size, geometry, and concentration of the susceptibility inclusions (Kiselev and Novikov, 2018; Yablonskiy and Haacke, 1994). For spherical susceptibility inclusions, the minimum radius above which the static dephasing regime is valid decreases with field strength as *B*_0_^−1/2^, and is thus lower at 11.7 T than at 3 T (Yablonskiy et al., 2021). We used the theoretical magnitude decay kernel, *a* = 1.26 kHz/ppm, for spherical inclusions in the static dephasing regime at 11.7 T (Chen et al., 2021; Yablonskiy and Haacke, 1994). However, the value of *a* may vary across different source geometries and in voxels where the static dephasing regime does not hold (Li et al., 2023). In the motional narrowing regime, the proportionality coefficient would theoretically depend on the diffusivity as well as the size and permeability of the susceptibility sources. For fitting the 3-pool model in DECOMPOSE-QSM, estimates of the nonlinear parameters (*χ*_*+,-*_) can be unreliable in voxels where the respective signal fractions (*C*_*+,-*_) are low (Chen et al., 2021). However, similar to the study by Chen et al., the PCS and DCS maps were observed to be robust with respect to the independent estimates of the nonlinear parameters. Lastly, while our study focused mostly on gray matter regions, future studies are needed to account for susceptibility anisotropy effects in white matter (Wharton and Bowtell, 2015).

## 5. CONCLUSION

In this study, we demonstrated the potential of quantitative susceptibility source separation to distinguish region-specific microgliosis and tau aggregation in the mouse brain. PCS and DCS maps showed significant regional differences between the WT and Tau mouse brains, that were found to correspond to localized microgliosis and phosphorylated-tau deposition, as validated by immunohistology. Intracellular tau accumulation and proliferation of microglia with iron loading are key pathological features of AD and other tauopathies. The results suggest that the paramagnetic and diamagnetic component susceptibilities may serve as sensitive imaging markers to detect these subcellular alterations, and may have useful applications to improve understanding of disease pathogenesis and to evaluate the efficacy of therapeutic treatments.

## Acknowledgements

This work was supported by the National Institutes of Health (NIH) under NIA grant R01AG057991, and NINDS grants R01NS124084 and R01NS127344.

## Notes

### Competing Interest Statement

The authors have declared no competing interest.

